# Higher cortical beta oscillatory power before balance perturbations is associated with worse balance in older adults with Parkinson’s Disease, but not in age-matched older adults

**DOI:** 10.64898/2026.09.02.748558

**Authors:** Andrew S. Monaghan, Jasmine L. Mirdamadi, Janna Protzak, Aiden Payne, Michael R. Borich, Lena H. Ting

## Abstract

Elevated beta activity is hypothesized to reflect reduced flexibility in updating motor and sensory states, which may impair the rapid integration of sensory information required for balance. Therefore, we predicted that higher cortical beta power immediately before balance perturbations would be associated with poorer clinical balance performance, particularly in individuals with Parkinson’s Disease (PD). Seventeen individuals with PD (median Hoehn & Yahr = 2) and eighteen neurotypical older adults stood on a translating platform while 32-channel electroencephalography was recorded. For each participant, the cortical component contributing most strongly to the perturbation-evoked N1 response, a balance-related cortical response localized to the supplementary motor area, was identified. Oscillatory beta power (13–30 Hz) during the one-second period immediately preceding perturbation onset was extracted from this N1-related cortical source and parameterized using Fitting Oscillations and One-Over-F (FOOOF). Balance ability was assessed using the Mini-Balance Evaluation Systems Test (Mini-BESTest). Beta power derived from this N1-related source did not differ between groups; however, within individuals with PD, higher pre-perturbation beta power was associated with lower Mini-BESTest scores, a relationship not observed in neurotypical older adults. A subset of older adults exhibited an additional, higher-frequency beta peak absent in PD, suggesting heterogeneity in cortical beta dynamics with aging. These findings suggest that elevated cortical beta oscillations immediately preceding balance perturbations are linked to clinically meaningful balance impairment in PD, supporting beta activity as a potential cortical biomarker of balance dysfunction in this population.

**New and Noteworthy:** This study provides novel evidence linking pre-perturbation beta oscillations over central sensorimotor regions to clinical balance impairments in people with Parkinson’s disease (PD), but not in neurotypical older adults. These findings highlight pre-perturbation beta activity as a correlate of balance deficits in PD and suggest its potential as a neural marker of balance impairment in this population.

## 1.0 Introduction

Balance impairment is a common and disabling symptom of Parkinson’s Disease (PD), yet the cortical mechanisms contributing to individual differences in balance function remain unclear. Sensorimotor beta oscillations (13-30 Hz) are elevated in both aging and PD (1–3) and may influence balance by constraining the flexible sensorimotor updating required for postural control. Beta oscillations are hypothesized to reflect interactions between cortical and subcortical motor regions (4), maintaining and modulating transitions between motor states (5, 6). In PD, abnormally strong beta oscillations, caused by disruptions in the basal ganglia-thalamocortical circuit due to dopamine loss, are linked to motor symptoms such as rigidity and bradykinesia (7–9). These pathological oscillations are thought to interfere with motor control by impairing normal beta suppression during movement, contributing to the motor deficits characteristic of PD (9, 10). Although beta power is higher in older adults, with and without PD, than in younger adults, the functional relevance of these oscillations for balance control remains unclear.

We hypothesize that elevated beta oscillations from the supplementary motor area (SMA) may impair balance by gating the sensory information needed to detect a balance disturbance and by constraining the motor flexibility needed to respond to it. Recovering balance after an unexpected perturbation depends on rapidly sensing how the body is moving and integrating this feedback into a corrective response. Beta oscillations are thought to gate this process, reducing the sensorimotor system’s sensitivity to incoming sensory input (6, 11); elevated beta may therefore blunt the feedback signaling that the body has been displaced, diminishing the ability to perceive and respond to a disturbance. Consistent with a role for beta in the perceptual aspects of balance, greater SMA beta activity is associated with poorer perception of body-motion direction in young adults (12), and in PD, impaired perception of whole-body motion is associated with worse balance ability, a relationship not observed in neurotypical older adults (13). Elevated sensorimotor beta may additionally constrain balance by resisting transitions between motor states, impeding shifts away from the current sensorimotor set, and slowing the motor updating required to execute a correction (6). However, it remains unknown whether the ongoing sensorimotor beta activity during standing balance relates to clinical balance ability, or whether this relationship differs between older adults with and without PD.

The balance N1, a perturbation-evoked cortical potential occurring approximately 100-200 ms after a balance disturbance, has been localized to the SMA (14–16), a region involved in integrating sensory information and motor planning during balance recovery (17, 18). This consistent and reliable event-related potential (19) can therefore be used to identify a participant-specific SMA source from EEG and measure that source’s activity during quiet standing. Inter-individual variability in the N1 tracks individual differences in balance and cognition across aging and PD: larger responses accompany poorer balance in young adults (20), whereas in older adults with and without PD, N1 amplitude, width, and timing relate to balance and cognitive function in group-specific ways (21–24). However, it remains unclear whether ongoing beta dynamics from the SMA prior to a balance disturbance contribute to these differences in balance performance between older adults and individuals with PD.

Here, we examined whether higher beta activity around the SMA during standing is associated with clinical balance ability in older adults with and without PD. Using spectral parameterization, we characterized both the oscillatory (periodic) and non-oscillatory (aperiodic) components of beta activity in the period immediately preceding balance perturbations, measured from a cortical source over the SMA. We then related these measures to clinical balance ability assessed with the Mini-BESTest, predicting that higher pre-stimulus beta power would be associated with poorer balance performance, particularly in individuals with PD.

## 2.0 Methods

### 2.1 Participants

Data from this cohort have previously been reported to examine the relationships between the balance-evoked N1 potential, cognitive function, and balance performance (25–27). The present study is a secondary analysis of previously reported participant data; analytic samples differ slightly across studies based on data availability and suitability for study-specific EEG analyses, including the use here of pre-perturbation EEG data that did not require identical perturbation parameters across participants. Seventeen older adults with PD (mean age 70 ± 7 years, median Hoehn & Yahr 2, 4 women) and eighteen older adults without PD (OA) (mean age 71 ± 7 years, 6 women) participated in the study. All participants provided written informed consent after receiving a thorough explanation of the experimental procedures, which were approved by the Emory University Institutional Review Board. Individuals with PD completed the experiment in the OFF-medication state, defined as being at least 12 hours after their last dose of dopaminergic medication. Each participant’s neurologist reviewed and approved an OFF-medication clearance before withholding medications for this study. Clinical and behavioral data were collected during the same OFF-medication session, and additional clinical records were consulted, when available, to verify disease duration and ensure participants met the inclusion criteria. Participants were recruited from the community surrounding Emory University and the Emory Movement Disorders clinic via flyers, outreach events, word-of-mouth, and databases of previous participants. Eligibility criteria for participants included being over the age of 55, having vision correctable to at least 20/40, no history of stroke or neurological conditions other than PD, no musculoskeletal issues causing pain or limiting leg mobility, the ability to stand unassisted for at least 15 minutes, and sufficient cognitive function to provide informed consent. Participants with prior experience on the perturbation platform, those on cholinergic medications, or those who did not receive neurologist approval to be OFF-medication were excluded.

### 2.2 Experimental Protocol

Participants stood on a platform that delivered 48 translational support surface perturbations to disturb standing balance (Figure 1A). The perturbations varied unpredictably in timing, direction (forward and backward), and magnitude (three levels) to challenge balance control and minimize adaptation. Each perturbation was followed by a quiet standing period of approximately 10 seconds before the next perturbation. EEG data were continuously recorded, and this study focused on the 1-second pre-stimulus period immediately preceding each perturbation (Figure 1A,B).

**Figure 1:**
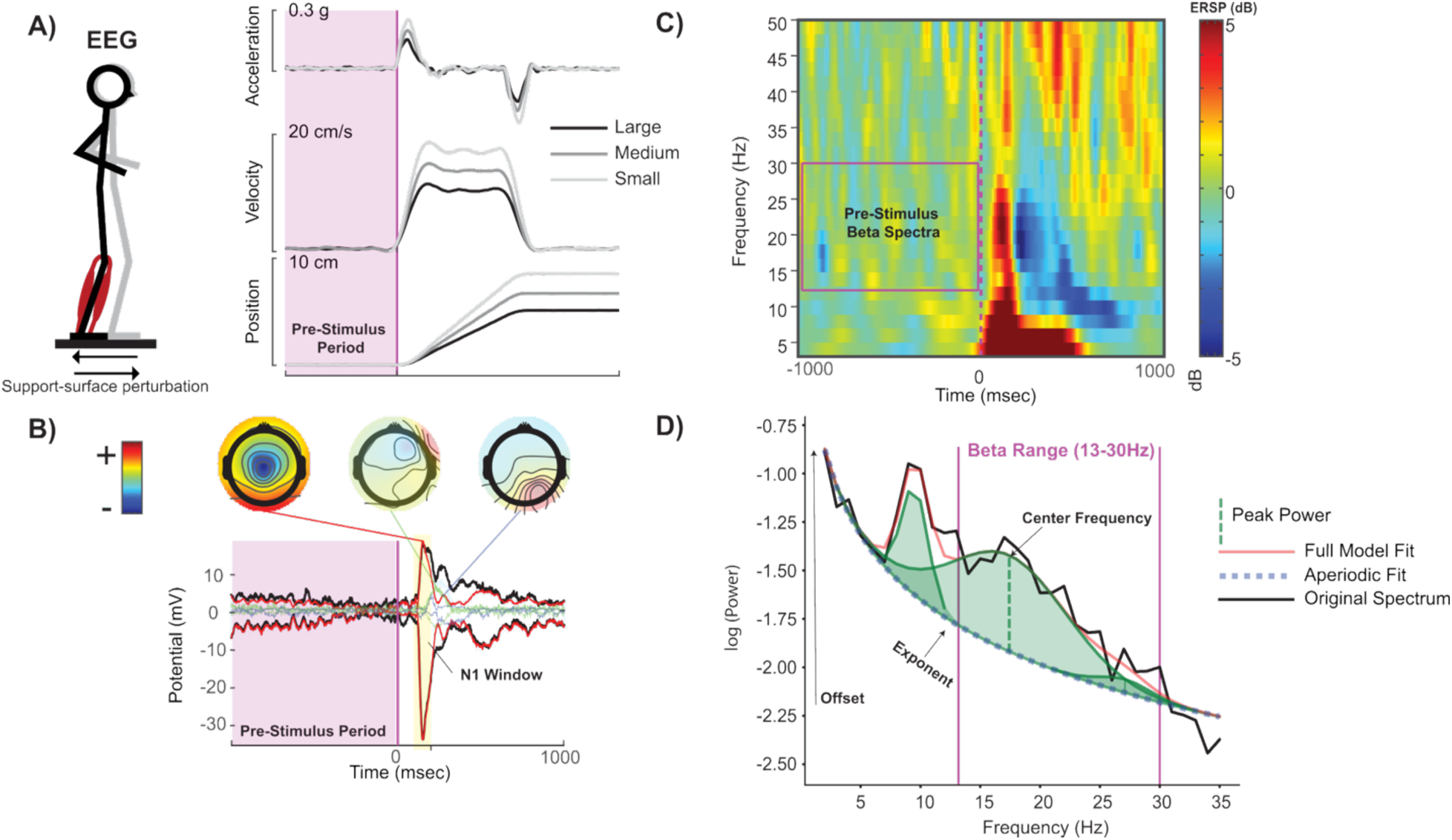
Experimental paradigm, EEG analysis, and spectral parameterization. (A) Participants experienced forward and backward support-surface perturbations at three intensity levels (small, medium, and large), while EEG was recorded. Only the pre-stimulus EEG (1 s before perturbation onset; magenta-shaded region) was used for analyses. Trials across all perturbation directions and intensities were averaged. (B) The largest source contributing to the N1 potential (100–200 ms; yellow-shaded region) was identified as the independent component (IC) explaining the highest percentage variance in this time window. (C) Pre-stimulus power spectra in the beta range (13–30 Hz) were derived from the N1-related IC. (D) Power spectra were parameterized into aperiodic (offset and slope) and periodic oscillatory components using the FOOOF algorithm.

### 2.3 Clinical Measurements

Balance performance was assessed using the Mini Balance Evaluation Systems Test (Mini-BESTest), which assesses four key domains: anticipatory postural adjustments, reactive postural responses, sensory orientation, and gait stability (28, 29). The test was scored according to the standard protocol, and for items requiring separate evaluations of the left and right sides, the lower score was recorded. Global cognition was assessed using the Montreal Cognitive Assessment (MoCA) (30), and the physical motor signs and symptoms of PD were evaluated with the Movement Disorder Society-Unified Parkinson’s Disease Rating Scale: Part III (MDS-UPDRS-III) (31).

### 2.4 EEG Recording and Preprocessing

EEG data were recorded using 32 active electrodes (ActiCAP, Brain Products, Germany) positioned according to the international 10–20 system, with two reference electrodes placed directly on the skin over the mastoids. EEG data were preprocessed using the EEGLAB toolbox in MATLAB (32). The data were downsampled from 1000 Hz to 500 Hz, and a 1 Hz high-pass filter was applied to remove low-frequency drift. The clean_rawdata plugin was used to identify and remove bad channels that were flat for more than 5 seconds, exhibited excessive high-frequency noise (>4 SDs), or were poorly correlated with neighboring channels (<0.6). Visual inspection verified these selections, and the remaining data were re-referenced to the average of all channels. To address power line noise, the Zapline-plus plugin was applied to remove 60 Hz interference (33, 34). Artifact Subspace Reconstruction (ASR) was used to correct non-stationary artifacts (35).

The data were epoched from −2 to 2 s around each perturbation onset and decomposed into maximally independent components (ICs) using the adaptive mixture independent component analysis (AMICA) algorithm (36, 37). ICs were categorized using the ICLabel plugin, an automated algorithm that classifies components as brain or non-brain sources (e.g., eye, muscle, and cardiac activity), and these classifications were confirmed with visual inspection (38). Non-brain ICs were removed from the dataset. Brain-related ICs were localized using the DIPFIT plugin in EEGLAB, which fits an equivalent current dipole model for each IC and maps them onto a standard Montreal Neurological Institute (MNI) template. ICs whose dipoles were located outside of the brain or that had a high residual variance (RV; >15%) between the observed scalp projection and the modeled dipole were excluded from further analysis (39, 40).

For each participant, the IC most strongly contributing to the N1 event-related potential (ERP), occurring 100–200 ms post-perturbation, was identified by calculating the percent variance accounted for (PVAF) within the N1 time window (100-200ms post-perturbation) using the pop_envtopo function in EEGLAB (Figure 1B). We selected this N1 IC for pre-stimulus beta analysis because it isolates the cortical source, presumed to be SMA, driving the balance-perturbation-evoked potential, making pre-perturbation beta oscillations more likely to reflect the sensorimotor activity relevant to perceptual-motor interactions for balance control (Figure 1B). If multiple ICs from a single participant contributed significantly to the N1 ERP, the IC with the largest percent variance accounted for was chosen (Figure 1B). For three participants, the N1-related ICs were manually selected based on the IC power spectrum and topographic map, guided by prior literature indicating that the N1 ERP is typically localized to the supplementary motor area (SMA) (14, 15).

### 2.5 Spectral Parameterization

The power spectral density (PSD) was computed for the IC contributing most to the N1 ERP (Figure 1C). For each participant, PSD calculations were performed on the pre-stimulus time window (-1000 ms to -2 ms relative to perturbation onset) using Welch’s method, with a Hamming window applied to 2-second segments and 50% overlap (Figure 1C). The spectral power was averaged across trials for each participant, collapsing across all perturbation directions (forward/backward) and magnitudes (small, medium, large). This approach was used since balance perturbations were delivered unpredictably in timing, direction, and magnitude, and the analysis focused on the quiet-standing period immediately preceding each perturbation, for which there was no a priori reason to expect systematic differences across upcoming perturbation conditions. The resulting PSD data were then analyzed using the FOOOF (Fitting Oscillations & One Over F) algorithm, which decomposes the power spectrum into periodic (oscillatory) components and the aperiodic (non-oscillatory) background activity (41) Figure 1D). This method allows characterization of neural oscillations by accounting for a 1/f-like aperiodic background, thereby isolating oscillatory peaks. The FOOOF parameters were set as follows: frequency range of 2–35 Hz, minimum peak height of 0, peak width limits of 1–12 Hz, a maximum of 4 peaks, a peak threshold of 2, and the aperiodic mode was fixed (Figure 1D).

Within the FOOOF-parameterized spectra, beta peaks were identified as local maxima in the 13–30 Hz range (Figure 1D). For each participant, we extracted the peak beta power (the power of the most prominent beta peak in the baseline window), average beta power (the mean of the peak-power values across all beta peaks detected within 13-30 Hz), peak beta center frequency (the frequency at which the most prominent beta peak occurred), and peak beta bandwidth (the width of the most prominent beta peak, reflecting the range of frequencies over which the beta oscillation occurs) (Figure 1D). Aperiodic parameters that provide insight into the non-oscillatory background of the neural signal were also estimated: the offset (the y-intercept of the aperiodic fit, representing overall signal power) and the exponent (the slope of the aperiodic component, reflecting the steepness of the 1/f-like background) (Figure 1D).

### 2.6 Statistical analyses

Differences in periodic and aperiodic metrics between PD and older adults were assessed using independent t-tests for normally distributed data. Effect sizes for t-tests were calculated using Cohen’s d (42), with thresholds of 0.20, 0.50, and 0.80 for small, medium, and large effects, respectively. Differences between groups with non-normal distributions were evaluated using Mann-Whitney U tests. Effect sizes for Mann–Whitney U tests were reported as r (Z/√N), which indexes the magnitude of the group difference rather than a linear association. Thresholds for r were 0.10, 0.30, and 0.50 for small, medium, and large effects, respectively (42). Pearson correlations assessed the association between beta power parameters and balance scores within each group. Multiple linear regressions examined the relationship between beta power parameters and balance performance across the PD and OA groups. Mini-BESTest scores, log-transformed to address non-normality and heteroscedasticity, served as the dependent variable. Independent variables included periodic and aperiodic beta power parameters, group (PD vs. OA), and their interaction terms. The mean-centering of beta power variables reduced multicollinearity and enhanced the interpretability of interaction effects (43). Model assumptions, including normality, homoscedasticity, and multicollinearity, were evaluated using Shapiro-Wilk tests, non-constant variance tests, and variance inflation factors (VIF), respectively. Influential points were identified using Cook’s distance (cutoff: 4/n) (44), and outliers were defined as observations with standardized residuals beyond ±2. Sensitivity analyses were conducted by refitting models after excluding influential points and outliers to assess the robustness of the findings (45). Effect sizes were reported using standardized regression coefficients (β) and partial eta-squared (η²ᵖ). For η²ᵖ, values of 0.01, 0.06, and 0.14 were interpreted as small, medium, and large effects, respectively (42). All statistical tests were two-tailed with a significance threshold of α = 0.05. Analyses were conducted using R version 4.0.3 (R Core Team, 2020).

## 3.0 Results

### 3.1. Group Characteristics

Demographic and clinical characteristics of the participants are presented in **Table 1**. There were no significant differences between the groups in age, height, weight, or cognitive function as measured by the Montreal Cognitive Assessment (MoCA). Both groups demonstrated relatively high cognitive function, as measured by the MoCA (OA: 26.2 ± 3.5; PD: 25.6 ± 2.7). However, individuals with PD had poorer balance performance than OA as measured by the Mini-BESTest (PD: 20.60 ± 6.30; OA: 24.60 ± 2.10; p = 0.021, d = 0.87). The PD group had a mean disease duration of 6.0 ± 3.2 years, with a median Hoehn & Yahr stage of 2 (IQR 2-3; n = 16), indicating moderate bilateral disease. Their motor symptom severity, as measured by the MDS-UPDRS III, averaged 30.6 ± 14.7 out of 132.

**Table 1:**
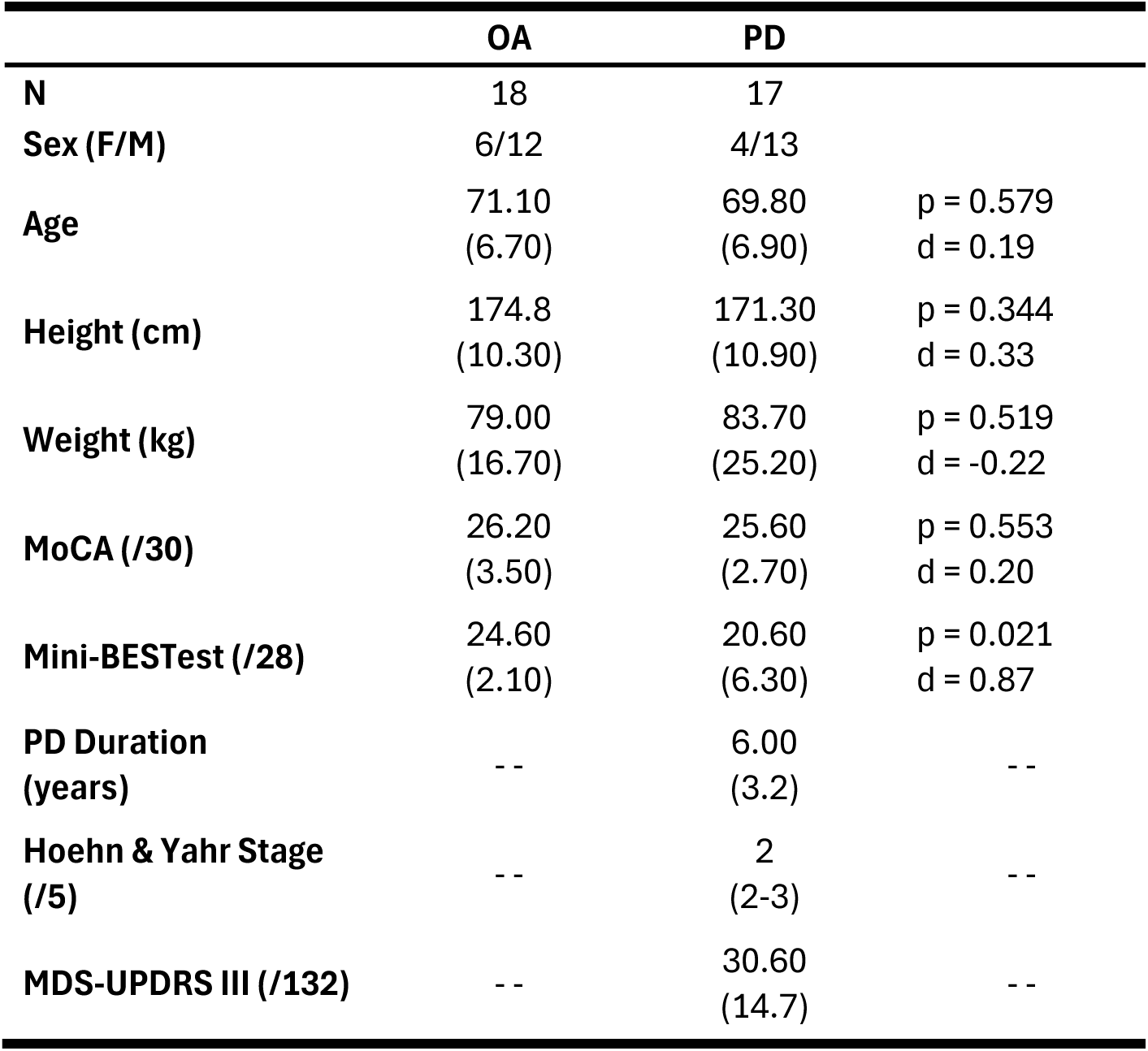
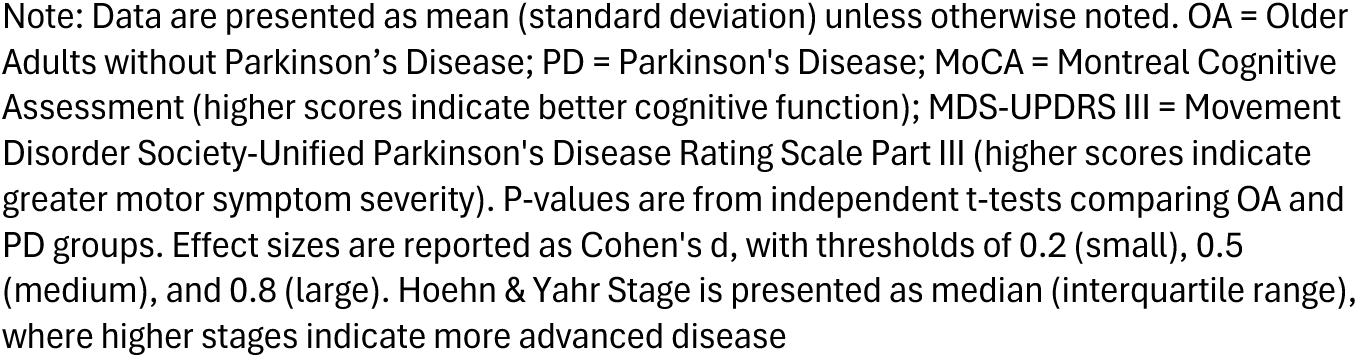
Demographic and Clinical Characteristics of Older Adults with and without Parkinson’s Disease.

|  | OA | PD |  |
| --- | --- | --- | --- |
| **N** | 18 | 17 |  |
| **Sex (F/M)** | 6/12 | 4/13 |  |
| **Age** | 71.10<br>(6.70) | 69.80<br>(6.90) | $p = 0.579$<br>$d = 0.19$ |
| **Height (cm)** | 174.8<br>(10.30) | 171.30<br>(10.90) | $p = 0.344$<br>$d = 0.33$ |
| **Weight (kg)** | 79.00<br>(16.70) | 83.70<br>(25.20) | $p = 0.519$<br>$d = -0.22$ |
| **MoCA (/30)** | 26.20<br>(3.50) | 25.60<br>(2.70) | $p = 0.553$<br>$d = 0.20$ |
| **Mini-BESTest (/28)** | 24.60<br>(2.10) | 20.60<br>(6.30) | $p = 0.021$<br>$d = 0.87$ |
| **PD Duration (years)** | -- | 6.00<br>(3.2) | -- |
| **Hoehn & Yahr Stage (/5)** | -- | 2<br>(2-3) | -- |
| **MDS-UPDRS III (/132)** | -- | 30.60<br>(14.7) | -- |

Note: Data are presented as mean (standard deviation) unless otherwise noted. OA = Older Adults without Parkinson's Disease; PD = Parkinson's Disease; MoCA = Montreal Cognitive Assessment (higher scores indicate better cognitive function); MDS-UPDRS III = Movement Disorder Society-Unified Parkinson's Disease Rating Scale Part III (higher scores indicate greater motor symptom severity). P-values are from independent t-tests comparing OA and PD groups. Effect sizes are reported as Cohen's d, with thresholds of 0.2 (small), 0.5 (medium), and 0.8 (large). Hoehn & Yahr Stage is presented as median (interquartile range), where higher stages indicate more advanced disease

### 3.2 N1 Component Selection and FOOOF Model Results

The N1 source accounted for most of the variance in the evoked response to support surface perturbations during standing balance. The primary ICs contributing to the N1 response were predominantly localized to sensorimotor areas, including the SMA (Supplemental Table 1). The mean percent variance accounted for was 85.61% (range: 46.69–98.13%) in the PD group and 91.85% (range: 49.25–100.00%) in the OA group. In the PD group (n = 17), 28 beta peaks were identified across participants, with one to three peaks per participant (median = 2). FOOOF model fits yielded R² values ranging from 0.761 to 0.993 (M = 0.960, SD = 0.050) and mean absolute errors (MAE) ranging from 0.030 to 0.107 (M = 0.056, SD = 0.018). In the OA group (n = 18), 37 beta peaks were identified, also with one to three peaks per participant (median = 2). Corresponding R² values ranged from 0.924 to 0.989 (M = 0.972, SD = 0.016), and MAE ranged from 0.033 to 0.090 (M = 0.052, SD = 0.015).

### 3.3 Group Comparisons of Beta Power Parameters Between PD and OA

Although beta power parameters did not differ significantly between PD and OA **(Figure 2**, **Table 2)**, a medium effect size indicated a trend toward higher peak beta center frequencies in older adults. No significant differences were observed in average beta power between the PD (M = 0.352, SD = 0.181 a.u.) and OA groups (M = 0.397, SD = 0.172 a.u.; p = 0.380, r = 0.15; **Figure 2C**, **Table 2**). Similarly, peak beta power did not differ significantly between individuals with PD (M = 0.372, SD = 0.178 a.u.) and older adults (M = 0.462, SD = 0.159 a.u.; p = 0.126, d = 0.53; **Figure 2D**, **Table 2**). Peak beta center frequency showed a non-significant trend (p = 0.051), with older adults (M = 20.18, SD = 3.91 Hz) tending to exhibit higher center frequencies than individuals with PD (M = 17.71, SD = 3.24 Hz; **Figure 2F**, **Table 2**). No significant differences were observed in peak beta bandwidth (p = 0.780, r = 0.05; **Figure 2E**, **Table 2**), aperiodic offset (p = 0.698, d = -0.13; **Figure 2G**, **Table 2**), or exponent (p = 0.738, d = 0.11; **Figure 2H**, **Table 2**).

**Figure 2.**
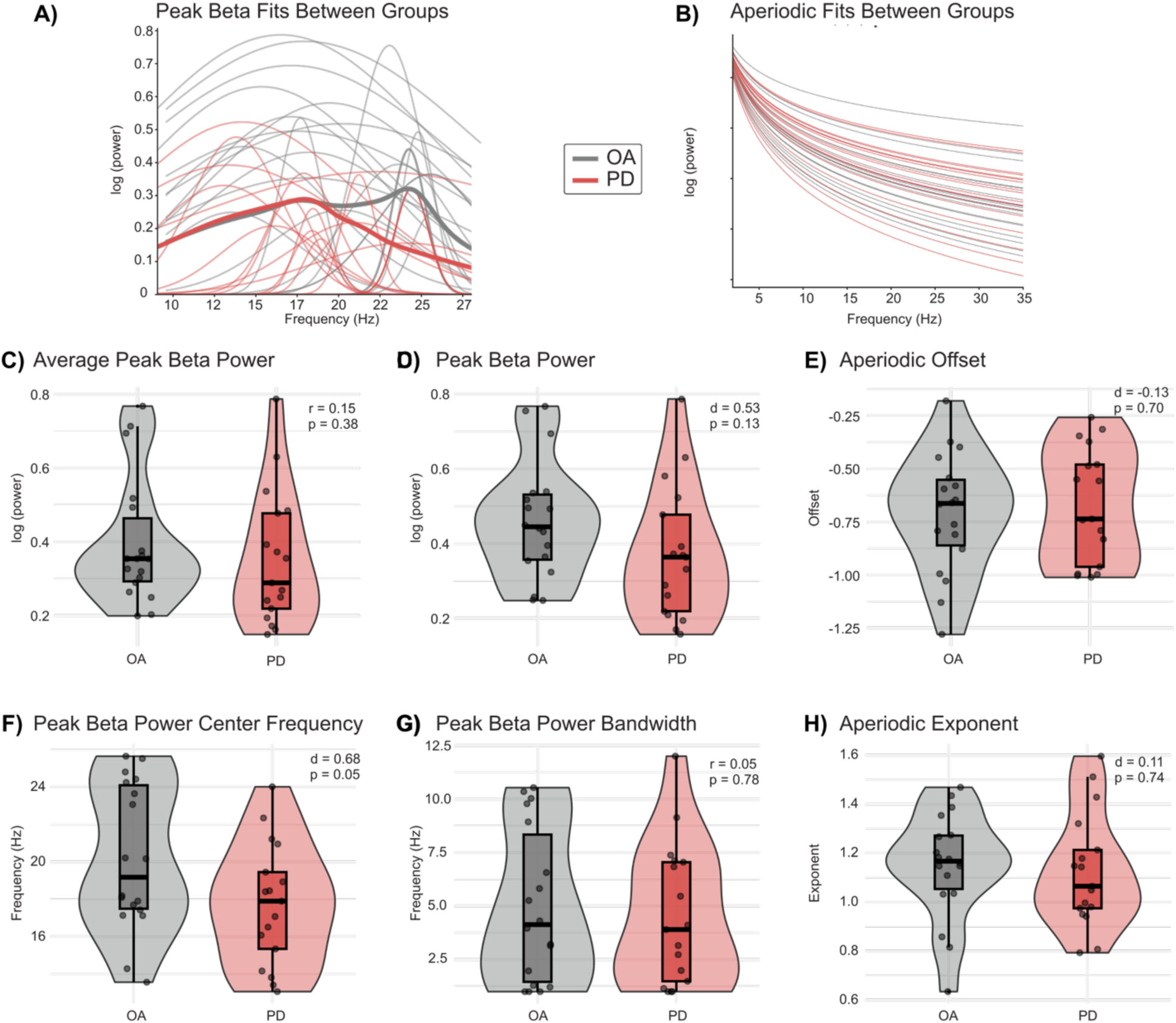
Group-level comparisons of beta power parameters in individuals with Parkinson’s disease (PD) and neurotypical older adults (OA). (A-B) Individual fits for periodic beta activity (A) and aperiodic parameters (B) across participants. (C-H) Violin and box plots comparing spectral metrics with individual data points overlaid. Statistical comparisons used α = 0.05, with effect sizes reported as Cohen’s d (small: 0.2, medium: 0.5, large: 0.8) and r (small: 0.1, medium: 0.3, large: 0.5).

**Table 2:** Beta Power Metrics in Older Adults with and without Parkinson’s Disease.

|  | OA | PD |  |
| --- | --- | --- | --- |
| N | 18 | 17 |  |
| Mean Beta Power | 0.397 (0.172) | 0.352 (0.181) | p = 0.380<br>r = 0.15 |
| Peak Beta Power | 0.462 (0.159) | 0.372 (0.178) | p = 0.126<br>d = 0.53 |
| Peak Beta Bandwidth | 4.962 (3.593) | 4.498 (3.303) | p = 0.780<br>r = 0.05 |
| Peak Beta Center Frequency | 20.176 (3.905) | 17.714 (3.238) | p = 0.051<br>d = 0.68 |
| Aperiodic Offset | -0.709 (0.283) | -0.673 (0.27) | p = 0.698<br>d = -0.13 |
| Aperiodic Exponent | 1.149 (0.218) | 1.123 (0.232) | p = 0.738<br>d = 0.11 |

**Note:** Data are presented as mean (SD). OA = older adults without Parkinson's Disease; PD = Parkinson's disease. Group differences were assessed using independent t-tests for normally distributed variables (effect size: Cohen's d) and Mann-Whitney U tests for non-normally distributed variables (effect size: r, calculated from the z-statistic). Avg. beta power represents the mean power across all detected beta peaks (13–30 Hz); peak beta power represents the largest beta peak. Peak beta center frequency indicates the frequency of the largest beta peak, and peak beta bandwidth reflects its width (Hz). Aperiodic offset and exponent are unitless parameters describing the 1/f background. Peak and average beta power and the aperiodic offset are expressed in $\log_{10}(\text{power})$ , in arbitrary units (a.u.), reflecting the FOOOF parameterization of independent-component power spectra.

### 3.4. Within-Group Associations and Interactions Between Beta Power Parameters and Mini-BESTest

In individuals with PD, greater beta power was associated with poorer balance performance. **Figure 3A** illustrates this relationship in exemplar PD participants spanning a range of balance ability, in whom lower balance ability corresponded to higher peak beta amplitude. Across the PD group, higher beta power was associated with lower Mini-BESTest scores (Figure 3B, red data points), with significant negative correlations for both mean (r = -0.67, p = 0.003) and peak beta power (r = -0.64, p = 0.006). In contrast, no significant associations were observed in the OA group (**Figure 3B, gray data points**; average beta power: r = -0.16, p = 0.52; peak beta power: r = -0.12, p = 0.64).

**Figure 3:**
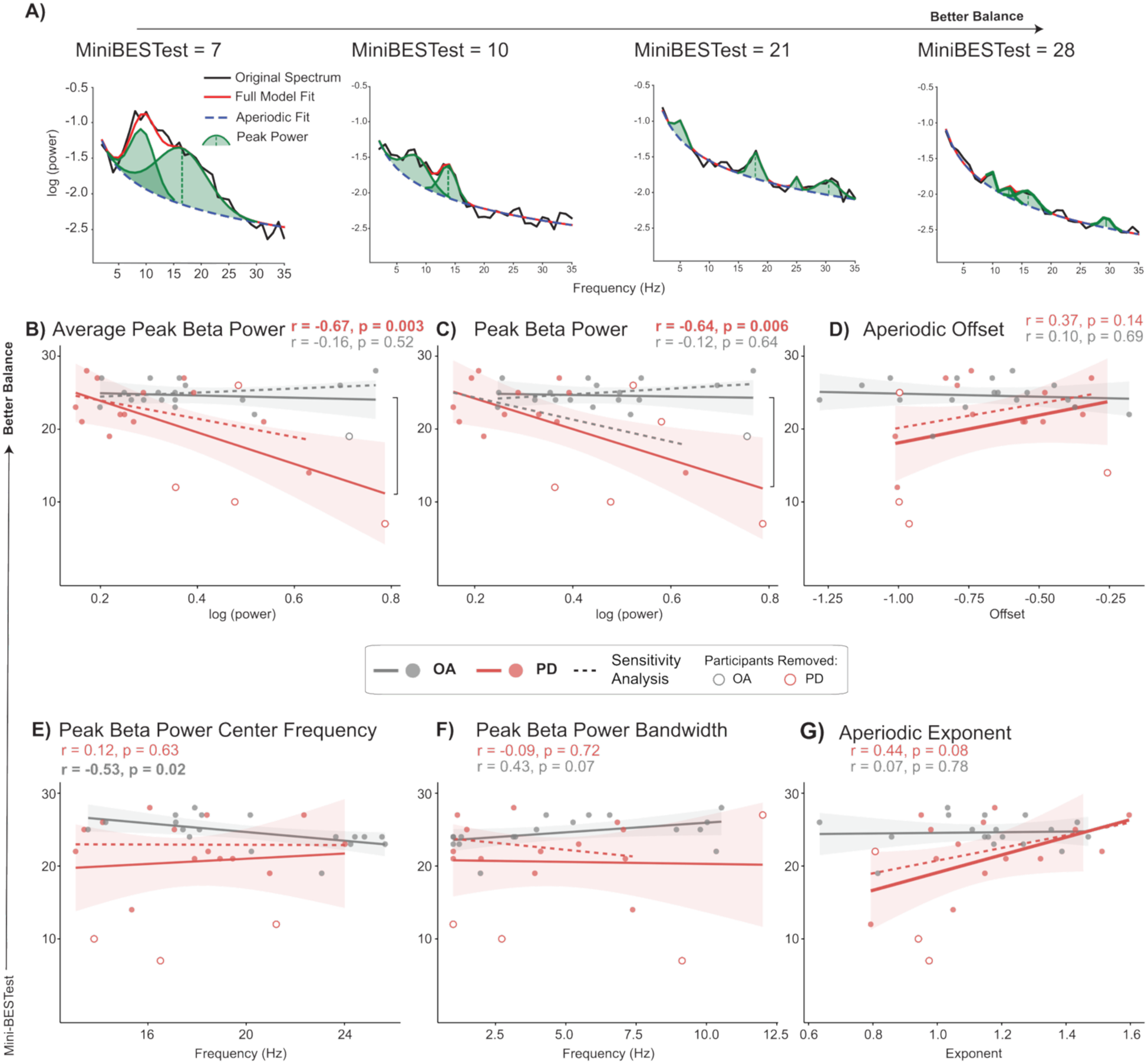
Higher beta power is associated with worse balance performance in Parkinson’s disease. (A) Representative power spectra and corresponding FOOOF model fits from PD participants spanning a range of balance performance (Mini-BESTest scores: 7–28). (B–G) Associations between Mini-BESTest scores and beta power parameters, shown separately for PD (red) and older adults without PD (OA; gray). Solid lines represent group-specific regression fits, with shaded regions indicating confidence intervals. Pearson correlation coefficients (r) are reported within each group. Significant group × beta parameter interactions are indicated by vertical brackets between regression slopes. Dashed lines represent sensitivity analyses excluding influential observations (Cook’s distance > 4/n) and outliers (standardized residuals ±2). Statistical significance was set at α = 0.05.

The relationship between beta power and balance performance differed between groups. Specifically, higher beta power was associated with poorer balance performance in PD but not in older adults (β = -1.37, 95% CI [-2.25, -0.49], p = 0.003, η²ᵖ = 0.25; **Table 3**) (Figure 3B, red filled circles vs. gray filled circles). This effect persisted after excluding influential observations in sensitivity analyses (**Figure 3B; red open circles and dashed regression lines, representing models refit after removal of influential observations; Table 3**).

**Table 3:**
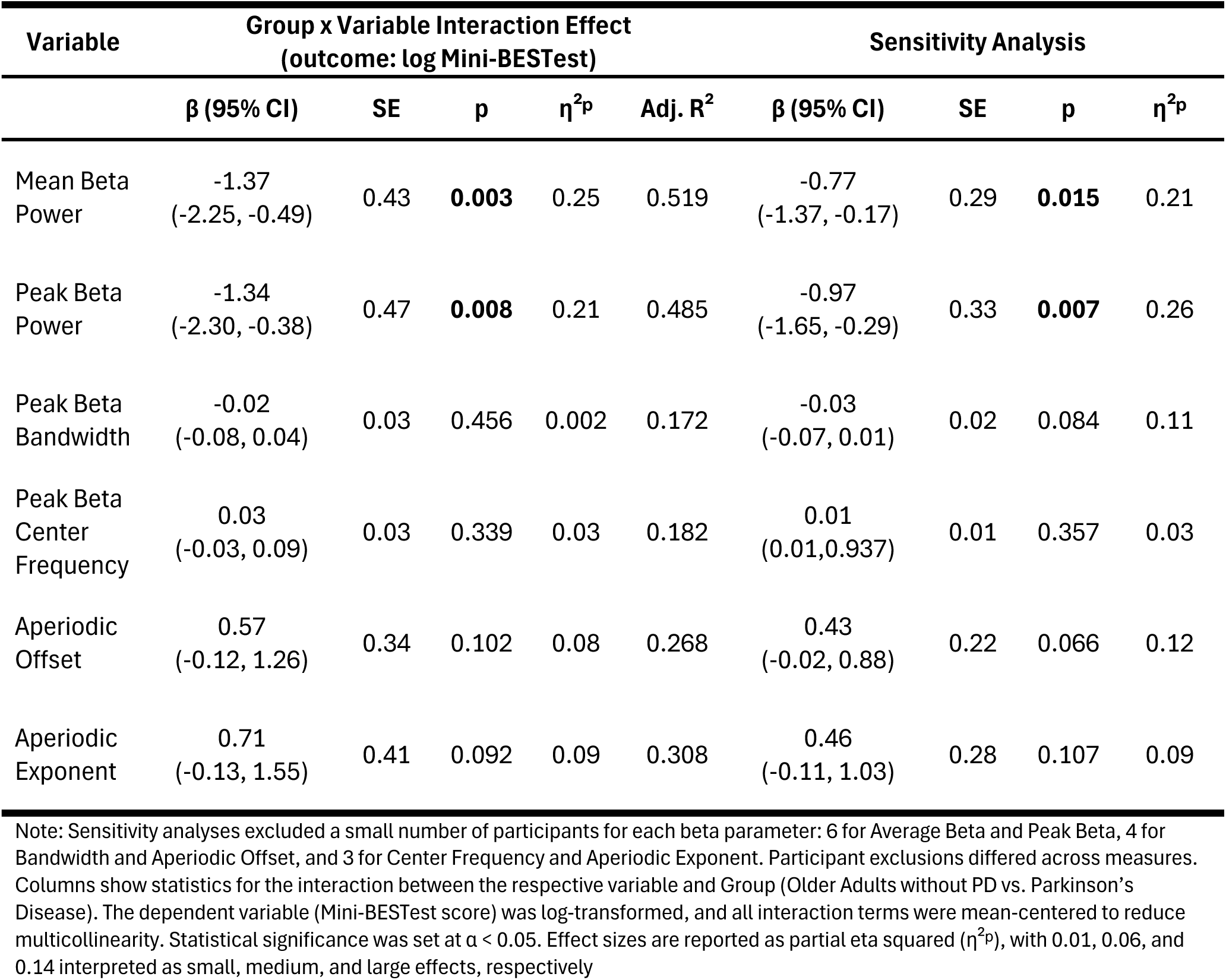
Beta Parameter x Group Interactions on Balance Performance (log Mini-BESTest) in PD and OA.

A similar interaction was observed for peak beta power (β = -1.34, 95% CI [-2.30, -0.38], p = 0.008, η²ᵖ = 0.21; **Table 3**), which also persisted after sensitivity adjustments (**Table 3**; **Figure 3C, red open circles and dashed regression lines**). While beta amplitude did not relate to balance in older adults (**Figure 3B-C, gray data points**), higher peak beta center frequencies were associated with lower Mini-BESTest scores in older adults (**Figure 3E, gray data points**; r = -0.53, p = 0.02), a relationship not observed in PD (**Figure 3E, red data points**; r = 0.12, p = 0.63). Peak beta power bandwidth (**Figure 3F**), aperiodic offset (**Figure 3D**), or aperiodic exponent (**Figure 3G**) showed no significant associations or interactions (**Table 3**).

## Discussion

Consistent with our prediction, higher SMA beta activity during standing was associated with poorer balance performance in individuals with PD, but not in older adults without PD, despite comparable beta power between groups. In PD, this association may reflect a constraint on the sensory and motor processing needed to detect and respond to a balance disturbance. In the older adult group, by contrast, beta varied considerably across individuals but was unrelated to balance, consistent with the more heterogeneous beta dynamics common to aging. These findings underscore the need to understand why elevated SMA beta activity relates to balance specifically in PD, and not in older adults with comparable beta power.

Our results suggest that higher pre-perturbation SMA beta activity may contribute to balance impairment in PD. This interpretation rests on the association between pre-perturbation SMA beta power and Mini-BESTest performance amongst individuals with PD, supporting the view that excessive beta activity constrains the sensory and motor processing required to detect and respond to a balance disturbance. Previous work provides evidence that cortical beta activity is altered during postural control in PD: individuals with PD show altered preparatory beta modulation before predictable perturbations, linked to reduced adaptability of postural responses (46), and perturbation-evoked cortical activity relates to balance ability, mobility, and motor symptom severity (25). This role for the SMA is anatomically grounded: as part of cortico–basal ganglia– thalamocortical circuitry, connected to the striatum and, via the hyperdirect pathway, the subthalamic nucleus (47, 48), the SMA is embedded in the dopamine-depleted networks whose elevated beta is linked to motor impairment in PD (49, 50), and its activity is altered with gait impairment and freezing (51, 52). Together, these functional and anatomical observations provide a plausible basis for a PD-specific relationship between pre-perturbation SMA beta and clinically relevant balance impairment.

The PD-specific brain-balance association may reflect constraints on the sensory processing needed to perceive a balance disturbance. Elevated beta may reduce the updating of sensory representations of body motion, lowering sensitivity to the incoming inputs required for accurate perception. Beta activity has been shown to modulate sensory sensitivity and perceptual processing (6, 11), and our prior work provides more direct evidence for this interpretation in the context of balance: greater pre-perturbation SMA beta activity was associated with poorer perception of whole-body motion in young adults (12). Consistent with the clinical relevance of this perceptual process, individuals with PD exhibit impaired whole-body directional acuity that is associated with poorer balance performance, a relationship not observed in older adults without PD (13). Together, these findings suggest that the association between higher pre-perturbation SMA beta and poorer balance observed here reflects, at least in part, reduced sensitivity to balance-relevant sensory information. Elevated beta could additionally constrain the motor updating required to respond to a disturbance, consistent with the proposed role of beta in maintaining the current sensorimotor state (6). However, because the present study related pre-perturbation beta to clinical balance ability rather than directly measuring perceptual or motor responses to the perturbation, the relative contributions of sensory and motor processes cannot be determined from the current data

In older adults without PD, beta variability appears to index heterogeneous individual differences rather than balance function specifically. Beta was not associated with clinical balance among older adults, consistent with prior work showing that neural measures relate differently to balance and cognition in PD and older adults (25). Specifically, older adults showed a trend toward higher peak beta center frequencies and a bimodal distribution of beta peaks. Higher beta center frequencies have been linked to motor and cognitive processes (5, 6) and may reflect broader individual differences in brain health or aging trajectories. Rather than reflecting a uniform shift with aging, these features may index differences such as cognitive reserve, physical activity, or overall brain health (53, 54), contributing to heterogeneity in beta dynamics that does not directly translate to balance ability. Consistent with the interpretation that age-related variability in neural dynamics reflects heterogeneous, dissociable underlying processes, prior work has shown that neural responses during balance in older adults relate to distinct, sometimes dissociable behavioral and cognitive factors rather than a single unified outcome (26, 55).

The absence of group differences in any beta parameter suggests that time-averaged beta power has limited utility for distinguishing PD from typical aging, and points instead to the temporal dynamics of beta as a potentially more informative marker. No oscillatory or aperiodic parameter differed between individuals with PD and older adults, consistent with prior work showing elevated cortical beta activity in both groups relative to younger adults, primarily measured during resting or seated conditions (2). Age-related increases in beta power have been observed across cortical regions using EEG and MEG in both resting and task-related contexts (1, 3, 56), while elevated beta activity in PD, including in the OFF-medication state, has been documented across cortical and subcortical motor circuits (57–59). This may reflect, in part, the nature of time-averaged spectral estimates. Beta activity in sensorimotor cortex occurs not as a continuous oscillation but in discrete transient bursts (4), and it is the rate and timing of these events, rather than their average amplitude, that appear most sensitive to disease-related changes. Vinding et al. demonstrated that PD patients OFF medication had lower cortical beta burst rates than healthy controls, while burst duration and amplitude were similar, and that lower burst rate correlated with worse bradykinesia and postural/kinetic tremor (60). Notably, beta burst rate has also been shown to increase with age in neurotypical adults (61), suggesting that burst dynamics may differentiate aging from PD even when mean power does not. Future work examining whether pre-perturbation beta burst characteristics similarly index balance impairment in PD may therefore provide greater sensitivity and specificity than the averaged power estimates used here.

## Conclusion and Future Directions

The present findings indicate that greater cortical beta power during standing derived from an SMA-related cortical source is associated with poorer balance performance in individuals with PD, but not in older adults without PD. These results suggest that, beyond serving as a potential marker of balance impairment in PD, greater beta may reflect a state of the sensorimotor cortex that constrains the perceptual and motor updating required for balance, although its spatial localization and mechanistic role remain to be fully established. Future longitudinal studies could clarify how changes in beta activity relate to balance decline and disease progression. In addition, experimental paradigms that independently manipulate sensory and motor demands may help determine whether beta activity in PD primarily reflects altered perceptual processing, motor control, or their interaction. Such approaches may clarify how ongoing sensorimotor beta activity contributes to the sensory and motor processes underlying balance impairment in PD, while further distinguishing disease-related from age-related beta dynamics

## Data and Code Availability

Data that support the findings of this study are available from the corresponding author (L.H.T.) upon reasonable request.

## Acknowledgements

We would like to thank our research participants. Study data were collected and managed using a Research Electronic Data Capture (REDCap) database hosted at Emory University.

## Author Contributions

A.P. and L.H.T. conceived and designed research; A.P. performed experiments; A.S.M. analyzed data; A.S.M, J.L.M., J.P, A.P., M.R.B., and L.H.T. interpreted results of experiments; A.S.M prepared figures; A.S.M drafted manuscript; A.S.M, J.L.M., J.P, A.P., M.R.B., and L.H.T. edited and revised manuscript; A.S.M., J.L.M., J.P., A.P., M.R.B., and L.H.T. approved final version of manuscript.

