## Supplemental Table 1 for "Higher cortical beta oscillatory power before balance perturbations is associated with worse balance in older adults with Parkinson’s Disease, but not in age-matched older adults"

**Supplemental Table 1 :** Characteristics of primary component contributing to the N1 potential for each participant

| Participant | X | Y | Z | PVAF | RV |
|---|---|---|---|---|---|
| ***OA*** | | | | | |
| 1 | -2.20 | -29.83 | 67.41 | 94.41 | 0.87 |
| 2 | -21.25 | -6.67 | 39.45 | 49.25 | 10.92 |
| 3 | -2.85 | -15.84 | 41.52 | 100.00 | 7.93 |
| 4 | -13.96 | -22.70 | 13.65 | 90.44 | 3.87 |
| 5 | 5.39 | -30.39 | 28.30 | 81.60 | 8.12 |
| 6 | 10.83 | -7.07 | 43.39 | 92.23 | 4.05 |
| 7 | 2.26 | -19.22 | 69.13 | 97.84 | 3.35 |
| 8 | -0.40 | -21.61 | 64.85 | 96.19 | 4.97 |
| 9 | 3.60 | -16.69 | 19.63 | 85.00 | 5.47 |
| 10 | 6.06 | 1.40 | 56.90 | 100.00 | 9.42 |
| 11 | -1.56 | -25.38 | 62.71 | 100.00 | 3.61 |
| 12 | 2.14 | -6.45 | 72.28 | 100.00 | 7.91 |
| 13 | 20.49 | -37.43 | 55.06 | 100.00 | 4.37 |
| 14 | -8.79 | -30.64 | 37.83 | 100.00 | 5.22 |
| 15 | 7.17 | -3.40 | 41.84 | 97.55 | 6.54 |
| 16 | -5.93 | -7.47 | 56.85 | 93.58 | 3.04 |
| 17 | -2.31 | -15.50 | 29.71 | 79.85 | 5.37 |
| 18 | 1.49 | -35.90 | 69.42 | 95.44 | 3.49 |
| ***PD*** | | | | | |
| 1 | 22.35 | -6.18 | 18.16 | 95.46 | 9.64 |
| 2 | 16.06 | -33.49 | 5.32 | 84.86 | 5.16 |
| 3 | -2.36 | -18.80 | 60.06 | 86.32 | 4.49 |
| 4 | 12.25 | -19.20 | 68.80 | 96.57 | 6.45 |
| 5 | 2.40 | -34.92 | 62.80 | 93.21 | 7.14 |
| 6 | 8.03 | -14.86 | 29.88 | 93.62 | 3.98 |
| 7 | -7.56 | -17.94 | 55.88 | 98.13 | 2.33 |
| 8 | -0.55 | -25.61 | 37.37 | 93.29 | 14.31 |
| 9 | 10.38 | -34.17 | 58.67 | 51.74 | 10.09 |
| 10 | 1.20 | -26.09 | 61.15 | 46.69 | 4.34 |
| 11 | -0.06 | -7.40 | 64.15 | 97.55 | 5.58 |
| 12 | -0.63 | -36.33 | 60.73 | 97.55 | 4.05 |
| 13 | 4.74 | -43.94 | 69.55 | 64.76 | 10.26 |
| 14 | -16.79 | -41.25 | 28.33 | 94.18 | 6.08 |
| 15 | 23.71 | -33.05 | 52.05 | 94.18 | 1.48 |
| 16 | 3.70 | -11.09 | 54.09 | 69.19 | 8.58 |
| 17 | 3.70 | -18.48 | 61.82 | 98.12 | 2.12 |

Note: X, Y, and Z coordinates represent the location of the equivalent dipole of the primary independent component (IC) in Montreal Neurological Institute (MNI) space. PVAF (Percent Variance Accounted For) indicates the proportion of the variance in the N1 event-related potential (100–200 ms post-perturbation) that the selected IC explains. RV (Residual variance) reflects the proportion of the variance in the scalp projection that is not accounted for by the dipole model.
